# IbinA and IbinB regulate the Toll pathway-mediated immune response in *Drosophila melanogaster*

**DOI:** 10.1101/2025.04.10.648149

**Authors:** Matthew K. Maasdorp, Susanna Valanne, Laura Vesala, Petra Vornanen, Elina Haukkavaara, Tea Tuomela, Aino Malin, Tiina S. Salminen, Dan Hultmark, Mika Rämet

**Author notes:** Corresponding author: Mika Rämet.

## Abstract

To combat infection, an immune system needs to be promptly activated but tightly controlled to avoid destructive effects on host tissues. IbinA and IbinB are related short peptides with robust expression upon microbial challenge in *Drosophila melanogaster*. Here, we show that *Ibin* genes are ubiquitously present in flies of the *Drosophila* subgenus *Sophophora*, where they replace a different but probably related gene, *Mibin*, which is found across a much wider range of cyclorrhaphan flies. Using synthetic peptides, we did not observe any direct bactericidal or bacteriostatic activity for either IbinA or IbinB *in vitro*. Using mutant *Drosophila* lines lacking the *IbinA* gene, *IbinB* gene, or both, we examined their roles in development and during microbial infections. *IbinA* is expressed in early pupae, and a lack of *IbinA* and *IbinB* leads to temperature-dependent formation of melanized tissue during metamorphosis, frequently around the trachea. IbinA and IbinB have distinct effects on susceptibility to microbial infection. For example, *IbinB* mutant flies, as well as flies lacking both *IbinA* and *IbinB*, had improved survival when challenged with *Listeria monocytogenes*, an intracellular pathogen, whereas a lack of *IbinA* alone had no effect. RNA sequencing of wildtype and mutant flies infected with *L. monocytogenes* showed enhanced Toll target gene expression in flies lacking *IbinB*, suggesting that IbinB acts as a negative regulator of the Toll pathway. In contrast, *IbinA* mutants had decreased Toll target gene expression in this context. Correspondingly, *IbinB* mutant flies had improved and *Ibin*A compromised survival in septic fungal infection, where the Toll pathway has a major role. Our study provides insight into the roles of IbinA and IbinB in regulation of the immune response in *Drosophila*.

**Author summary:** While the immune systems of animals must be able to be rapidly activated, they have the potential to cause severe tissue damage when overactive. *Drosophila melanogaster* has proven to be a highly effective model for studying core immune system pathways, and for developing concepts in host-pathogen interactions. The main signaling pathways involved in activation of the *Drosophila* immune system are well described and have contributed to the discovery of homologous pathways in the mammalian immune system. Despite this, the molecular mechanisms of action of the genes expressed downstream of these pathways are generally not well characterized. Regulation of the immune response also requires further investigation. Here, we use *Drosophila* mutants to show roles for two short peptides, both highly expressed during infection. IbinA and IbinB regulate the humoral immune response and the melanization reaction (an insect-specific immune reaction against parasites). Flies lacking one or both of these genes mount a stronger, more effective response to some pathogenic bacteria compared to wild type flies. During development, IbinA and IbinB have tissue protective roles, with mutants showing tissue damage due to aberrant immune activation. Our results indicate that IbinA and IbinB are important regulators of the immune response with tissue protective properties.

## Introduction

*Drosophila melanogaster* has an elegant innate immune system that includes both cellular and humoral arms (1). The cellular immune response involves mechanisms such as pathogen recognition, phagocytosis of bacteria, and encapsulation and the killing of parasitoid wasps (2–5), and is mediated by *Drosophila* immune cells called hemocytes. The *Drosophila* humoral immune response is initiated by the recognition of Pattern Associated Molecular Patterns (PAMPs) by Pattern Recognition Receptors (PRRs), resulting in the release and nuclear localization of NF-κB transcription factors (Dif, Dorsal, and Relish) via the Toll and Immune deficiency (Imd) signaling pathways, which are conserved between flies and humans (6–10). Whereas the microbial recognition mechanisms and the core signaling pathways leading to activation of the humoral immune response via the Toll and the Imd pathways have been well established, the immune effector mechanisms are largely yet to be characterized at the molecular level. Some of the induced genes encode secreted peptides with direct antimicrobial properties, like Cecropins, Attacins, Diptericins, insect Defensins, Drs and Mtk (11–16), whereas some, like *pirk*, are involved in the regulation of the immune response (17–20). The role of many infection-inducible genes is, however, currently unknown.

In our previous work, we identified novel immune-induced molecules, namely Induced By INfection (IBIN, CG44404 [CR44404]) and CG45045 (CR45045) that are rapidly induced in immune challenge via both the Toll and the Imd pathways (21). It was originally thought that IBIN and CG45045 are non-coding RNA molecules, but they have been re-annotated as peptide-encoding genes with strong sequence similarity to each other (21,22). Because of their similarity, we have renamed the peptides as IbinA (IBIN, CG44404) and IbinB (CG45045). The *IbinA* gene is predicted to encode a 41 amino acid-long peptide, whereas the *IbinB* gene gives rise to a putative 42 amino acid-long peptide homologous to IbinA. *IbinA* and *IbinB* are highly upregulated by both gram-positive and gram-negative bacteria, as well as when fly larvae are parasitized by *Leptopilina boulardi* wasps (21). Upon infection, *IbinA* is expressed in the fat body (the functional equivalent of mammalian liver and adipose tissue, and an important immune tissue in the fly), in hemocytes (fly blood cells), and in the gut. Interestingly, ectopic *IbinA* expression in the fat body and in the gut alters expression of sugar metabolism genes and downregulates protein metabolism genes (21). In addition, others have shown that *IbinA* is induced in female flies upon sight of parasitoid wasps (23), and in male flies upon social isolation (24), suggesting a role for Ibin peptides also in other stress-related situations besides infection. Here, we set out to investigate the roles of IbinA and IbinB in *Drosophila* immunity using deletion mutant lines for the *IbinA* and *IbinB* genes, as well as a double mutant line lacking both genes.

## Results

### *IbinA* and *IbinB* are part of a family of genes conserved across fly species

We identified *Ibin* homologs by blastp and tblastn searches in many *Drosophila* species (examples are given in Fig 1A and S1 Fig), but only within the subgenus *Sophophora* (including *Lordiphosa* and the willistoni and saltans groups). To get insight into the origin of this apparent novelty, we looked at the corresponding chromosomal regions in other fly species. In *D. melanogaster* the *IbinA* gene is located between the *CG30109* and *p32* genes in chromosome 2. In terms of the wider chromosomal context, Ibin genes are located near the Diptericin/Attacin loci (S2 Fig). In species that lack *Ibin* homologs, the corresponding chromosomal locus encodes a proline-rich peptide, which we call ‘Mother of Ibin’ (Mibin, S2 Fig).

**Figure 1.**
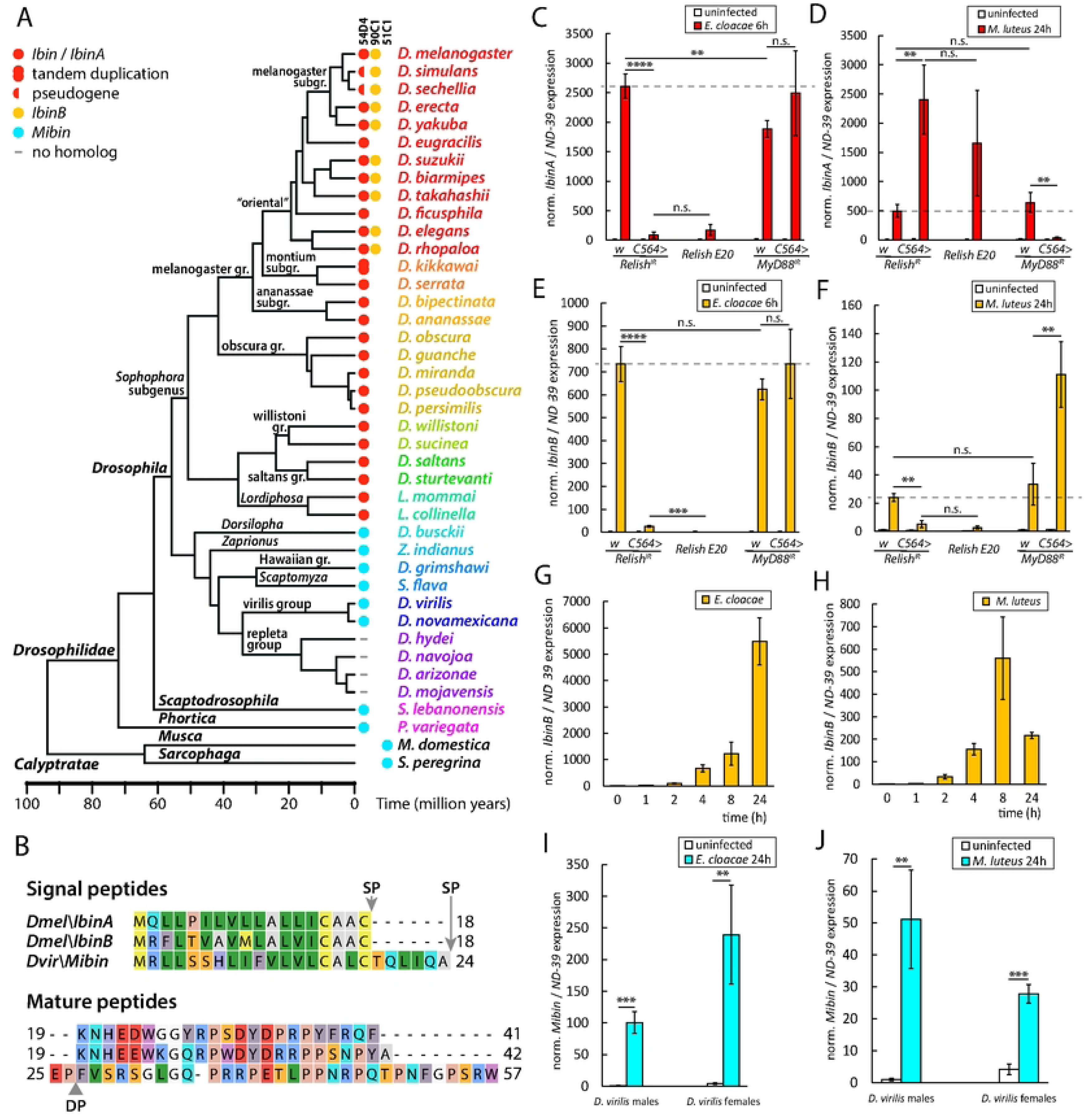
*IbinA* and *IbinB* belong to a family of genes conserved across fly species. **A)** Consensus phylogenetic tree showing the occurrence and chromosomal locations (54D4, 90C1 or 51C1) of *IbinA, IbinB* and *Mibin* orthologs in representative drosophilid flies. Time calibration is based on (26), the calyptrate outgroup is not time calibrated. **B)** Predicted IbinA and IbinB peptide sequences with the predicted cleavage sites for signal peptide (SP, likelihood: 0.998) marked by arrows and for dipeptidyl peptidase (DP) marked by a triangle. *IbinA* or *IbinB* expression in *D. melanogaster* when challenged **C)** and **E)** with *E. cloacae* or **D)** and **F)** with *M. luteus*. **G-H)** Expression patterns of *IbinB* measured within the first 24 hours after *D. melanogaster* exposure to *E. cloacae* or *M. luteus*. **I-J)** *Mibin* expression in *Drosophila virilis* males and females when exposed to *E. cloacae* or *M. luteus*. **C-J)** Statistically significant differences are marked with asterisks: * = p<0.05; ** = p<0.01; *** = p<0.001, **** = p<0.0001. For fold induction values, expression values in uninfected flies were set to 1.

Conserved *Mibin* genes could be identified among most *Drosophilidae* as well as in other acalyptrate and calyptrate flies, like the house fly (*Musca domestica*) and the flesh fly (*Sarcophaga peregrina*), and even in aschizan flies, like the hoverflies, family Syrphidae (S3 Fig). Thus, the *Mibin* genes must go back at least to the origin of the first cyclorrhaphan flies, perhaps about 110 million years ago (25). Importantly, we could not find *Mibin* homologs in any of the species that have *Ibin* genes (Fig 1A), which led us to hypothesize that the original *Ibin* gene has evolved directly from *Mibin* in an early ancestor of the subgenus *Sophophora*, about 55 million years ago (26). Later, a duplication of the ancestral *Ibin* gene produced the paralogs *IbinA* and *IbinB*, of which only *IbinA* remains in the original location at 54D4, while *IbinB* has translocated to 90C1 (chromosome 3). We identified separate *IbinA* and *IbinB* genes in most members of the *Drosophila* oriental subgroups, but not in other subgroups (Fig 1A), suggesting that the duplication happened in an ancestor of the oriental subgroups, probably less than 30 million years ago (Fig 1A). It could also have happened earlier if one copy was secondarily lost e.g., in the montium and other subgroups.

Both IbinA and IbinB are predicted with a high likelihood (0.998) to have a signal sequence at the N-terminus. The signal sequence is predicted to be cleaved between amino acids at positions 18-19, with a probability of 0.957 for IbinA and 0.951 for IbinB (Fig 1B). After cleavage, the length of the predicted mature IbinA peptide is 23 amino acids; the IbinB peptide is 24 amino acids long. Mibins also have predicted signal sequences for export, but these peptides may also be subject to additional processing. In these peptides, the signal peptidase cleavage site is often followed by one or more dipeptides of the general format Xxx-Pro (XP), which are potential substrates for dipeptidyl peptidase IV-like enzymes (Fig 1B). Of 144 investigated Mibin sequences, 102 had at least one N-terminal XP dipeptide (most commonly EP), and 13 of them had tandem repeats of 2-3 XP dipeptides (S3 Fig). Similar amino-terminal dipeptides have been found in many insect antimicrobial peptides, like cecropin, diptericin, drosocin and hymenoptaecin, as well as in insect venom peptides (27–32). For cecropin and the bee venom melittin, the N-terminal dipeptides have to be removed in order to activate the peptides (27,33). Despite their proposed relationship, the mature Mibins have no significant sequence similarity to IbinA or IbinB, beyond the fact that several proline and glycine residues are present in each of these peptides (Fig 1B). The Ibin family members are characterized by a conserved KNHEEWXG motif at the N-terminus (S1 Fig). By contrast, the Mibin peptides share a C-terminal proline-rich motif, RPXTLPPN/QRPXXPN/DF (S3 Fig), which has no counterpart in the Ibin peptides.

*IbinA* expression was previously shown to be strongly induced by bacterial infection in response to the Imd and Toll signaling pathways (21). We have now extended these studies and compared the responses of *IbinA* and *IbinB* expression in selected conditions by RT-qPCR. We assayed the response to *Enterobacter cloacae*, which is a strong inducer of the Imd pathway, and *Micrococcus luteus*, which activates Toll signaling. The response was studied in *D. melanogaster* wildtype flies compared to *C564>Relish^IR^*, or *Relish^E20^* null mutant flies (with impaired Imd pathway signaling), or to *C564>MyD88^IR^* flies with *MyD88* knockdown in the fat body (impaired Toll pathway signaling). Expression of both *IbinA* and *IbinB* is induced by both bacteria, but the inducibility of *IbinA* (Fig 1C-D) (compared to baseline) is stronger than that of *IbinB* (Fig 1E-F), especially with *M. luteus* infection (Fig 1D & 1F). Incidentally, we noticed that *M. luteus* infection causes a small decrease in the expression of the selected control gene *ND-39* compared to the uninfected flies. This effect was taken account while analyzing the results. As also shown previously (21), *IbinA* expression induced with *E. cloacae* is dependent on *Relish*, and thus on the Imd pathway (Fig 1C), whereas when infected with *M. luteus*, *IbinA* expression depended on the *MyD88* gene in the Toll pathway (Fig 1D). Of note, like *IbinA*, *IbinB* expression was dependent on the Imd pathway when induced with *E. cloacae* (Fig 1E), but, unlike *IbinA* expression, *M. luteus*-induced *IbinB* expression was also dependent on the expression of Imd pathway gene *Relish* (Fig 1F). To study this further, we analyzed the importance of known Imd pathway genes (*PGRP-LC*, *Imd* and *FADD*) on *IbinB* expression upon *M. luteus* challenge. Silencing these genes blocked the induced *IbinB* expression (S4A, S4B Fig) further supporting the idea that the Imd pathway is needed for both *E. cloacae*-and *M. luteus*-induced *IbinB* expression.

Expression kinetics of *IbinB* (Fig 1G) resemble those of *IbinA* after exposure to *E. cloacae*, whereas expression of *IbinB* peaks earlier after *M. luteus* infection (Fig 1H) than seen with *IbinA* (21). *Drosophila virilis Mibin* expression was also strongly induced by the abovementioned bacteria (Fig 1I-J), giving further support to the relatedness of the *Ibin* and *Mibin* genes. These data indicate that *IbinA* and *IbinB* are conserved among *Drosophila* species and their expression is controlled by NF-κB signaling upon microbial challenge, suggesting a role in innate immunity.

### *IbinA* shows variable expression in aseptic stress conditions

Previous studies have shown upregulation of the *IbinA* gene, together with a handful of antimicrobial peptide and other immune-responsive genes, in the heads of female flies when they are visually exposed to parasitoid wasps (23), and in the heads of male flies when they are socially isolated (24). This suggests that IbinA and IbinB might have stress-related functions. To study this hypothesis further, we first performed RT-qPCR from heads of flies subjected to the abovementioned conditions. This led to a similar *IbinA* induction as published earlier (S4C Fig). This expression was highly variable between samples, and, at a much lower level compared to that seen in infected flies. We also attempted to replicate the behavioral findings of Ebrahim and coworkers (23) by expressing *IbinA* under the control of the pan-neuronal *elav-GAL4* driver in the female flies. However, this overexpression did not result in delay to mating (S4D Fig). As previous work has focused on *IbinA* expression, we looked at whether *IbinB* is also upregulated in the fly head in this situation. We did not see a significant increase in *IbinB* expression when female flies were visually exposed to wasps (S4E Fig). Similarly to the wasp exposure results, single-housed (socially isolated) male flies show a moderate increase in expression of *IbinA* in their heads (S4F Fig), replicating the results from Agrawal and colleagues (24). To further define the situations in which *IbinA* is expressed, we studied the expression patterns of these genes under other physiological stress conditions. *IbinA* was not significantly upregulated by heat shock (S4G Fig), or osmotic stress (S4H Fig). These data indicate that infection is the predominant inducer of *IbinA* and *IbinB*.

### IbinA and IbinB do not have bactericidal or bacteriostatic properties

While *IbinA* and *IbinB* genes are immune-inducible and the (predicted) peptides are well conserved across related species, their role in fly immunity remains unknown. As short peptides that are produced minimally in uninfected flies, and highly induced upon infection (Fig 1C-F), IbinA and IbinB bear resemblance to AMPs and might have a function in resistance against invading pathogens. To test if IbinA and/or IbinB can prevent bacterial reproduction or directly kill bacteria, we incubated two enterobacteria (*E. cloacae* and *Escherichia coli*) and two cocci (*Staphylococcus aureus* and *Enterococcus faecalis*) with synthetic, mature IbinA and IbinB peptides. The antimicrobial peptide Cecropin A was used as a positive control for the enterobacteria, whereas Melittin (from bee venom) was used as a positive control for *S. aureus,* and Lysozyme (from egg white) for *E. faecalis* (Fig 2). We did not find any effect of IbinA or IbinB peptides on the bacterial growth at any concentration, in contrast to the positive controls, which were able to clear bacteria in a concentration-dependent manner (Fig 2). These data suggest that although IbinA and IbinB are likely to have an immune response-related function, they do not act as antimicrobial peptides, at least not in isolation.

**Figure 2.**
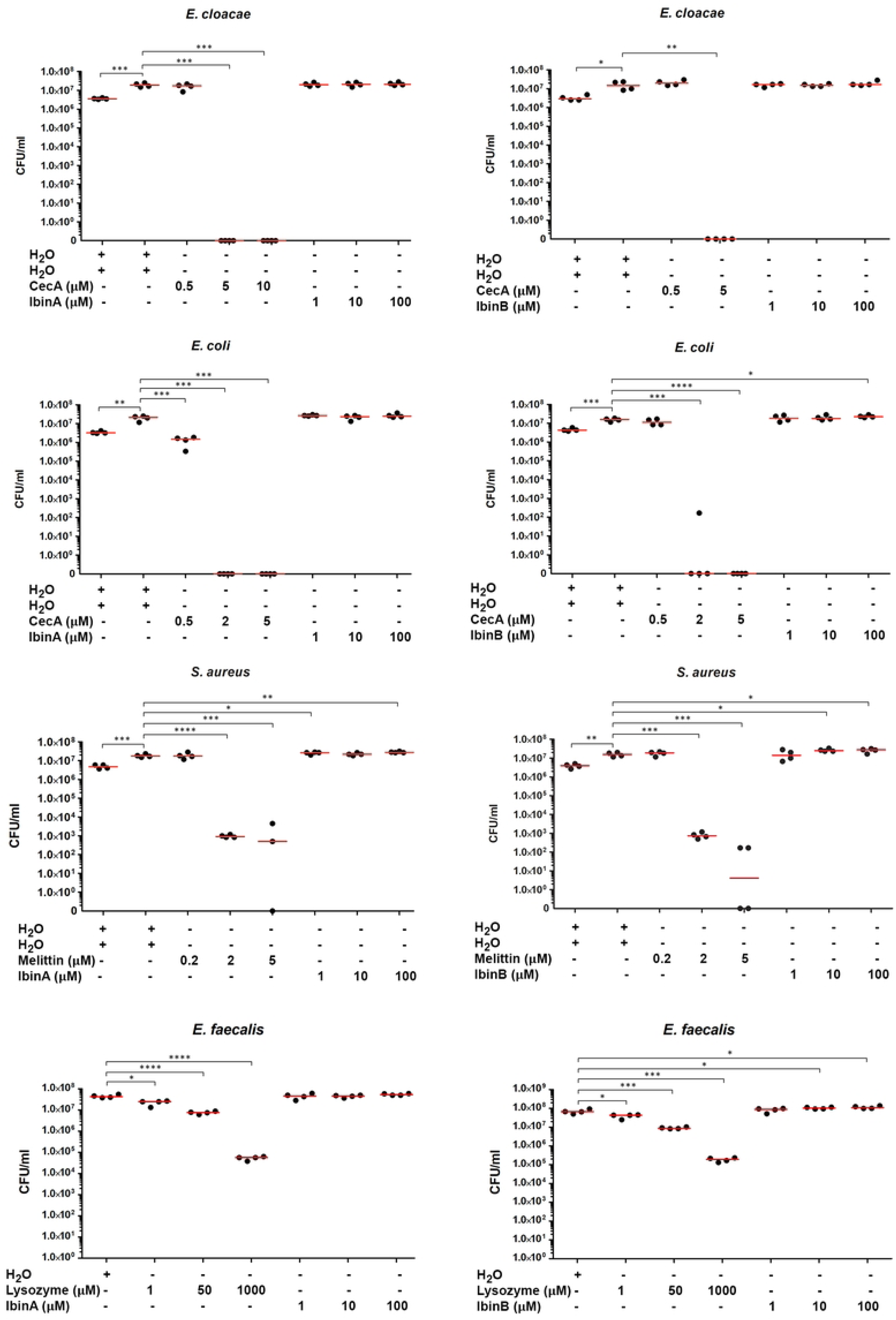
IbinA and IbinB synthetic peptides do not kill bacteria *in vitro*. The following positive controls were used: Cecropin A for *E. cloacae* and *E. coli*, Melittin for *S. aureus* and Lysozyme for *E. faecalis*. Statistically significant differences are marked with asterisks: * = p<0.05; ** = p<0.01; *** = p<0.001, **** = p<0.0001.

### *IbinA* and *IbinB* mutant flies exhibit developmental and blood cell phenotypes and changes in lifespan

To characterize the biological roles of IbinA and IbinB, we had knock-out mutant fly lines for the *IbinA* and *IbinB* genes generated. Precise deletions of *IbinA* (*IbinA^KO^*) and *IbinB* (*IbinB^KO^*) genes were generated using Crispr/Cas9 technology while simultaneously knocking in a selection marker as detailed in materials and methods and schematically illustrated in Fig 3A & B. Furthermore, a double mutant line for both *IbinA* and *IbinB* (*IbinA^KO^*; *IbinB^KO^*) was produced by crossing the single mutant lines together. *IbinA* was not expressed in *E. cloacae*-infected *IbinA^KO^* mutant or in the double mutant (*IbinA^KO^*; *IbinB^KO^*) (Fig 3C) and similarly, *IbinB* was not expressed in the *E. cloacae*-infected *IbinB^KO^* mutant or in the double mutant (Fig 3D). Overall lifespan showed significant differences between mutant and wild-type lines, with *IbinA^KO^* flies showing longer lifespan, and *IbinB^KO^* and double mutant showing shorter lifespan (Fig 3E). Some of the double mutant flies were also noted to have areas of melanized tissue visible through the cuticle (Fig 3F_iii_-3F_vi_: example images, compare to Fig 3F_i_-3F_ii_ wildtype flies lacking abnormal melanization, 3G: quantification). These areas of melanization frequently occurred in the abdomen at locations of spiracles and trachea (examples in Fig. 3F_v_ and 3F_vi_, yellow arrowheads), while some individuals (both those with and without the larger spiracle/tracheal melanization spots) showed smaller melanized spots visible through the dorsal cuticle (Fig. 3Fiii and 3Fiv, red arrow heads). Melanized tissue was seen in *IbinA^KO^* and *IbinB^KO^* single mutant flies much less frequently. Penetrance of this phenotype varied, with higher numbers of flies exhibiting melanized tissue when reared at 25°C compared to 18°C (Fig 3G). The melanization phenotype was also found at higher rates in females compared to males (especially in *IbinA^KO^;IbinB^KO^* flies at 25°C). At 29°C mutant flies developed normally until emergence from the pupal case, at which point only 11% of double mutant flies eclosed, while *IbinA^KO^* and *IbinB^KO^* flies eclosed at much higher rates, occasionally exhibiting small areas of melanization in points under the dorsal cuticle (Fig 3G). This was seen most often in *IbinA^KO^* flies, again, particularly in females. We did not observe melanized tissue in larvae, but this phenotype was present in some individuals already at pupal stages (example images Fig 3H, compare wildtype in Fig 3H_i_ to individual with melanized tissue in Fig 3H_ii_). Flies with large areas of melanized tissue were sometimes observed to have brown discoloration on the dorsal side of the abdomen, putatively pericardial cells taking up melanized material (S5A_i_ Fig, purple arrowhead). We also observed that male *IbinA^KO^* flies frequently have areas missing from the normally expected bands of coloration on the abdomen (S5A_ii_ Fig, green arrowhead), similar to the phenotype observed by Scherfer and coworkers when they knocked down *Serpin-28D* (34).

**Figure 3.**
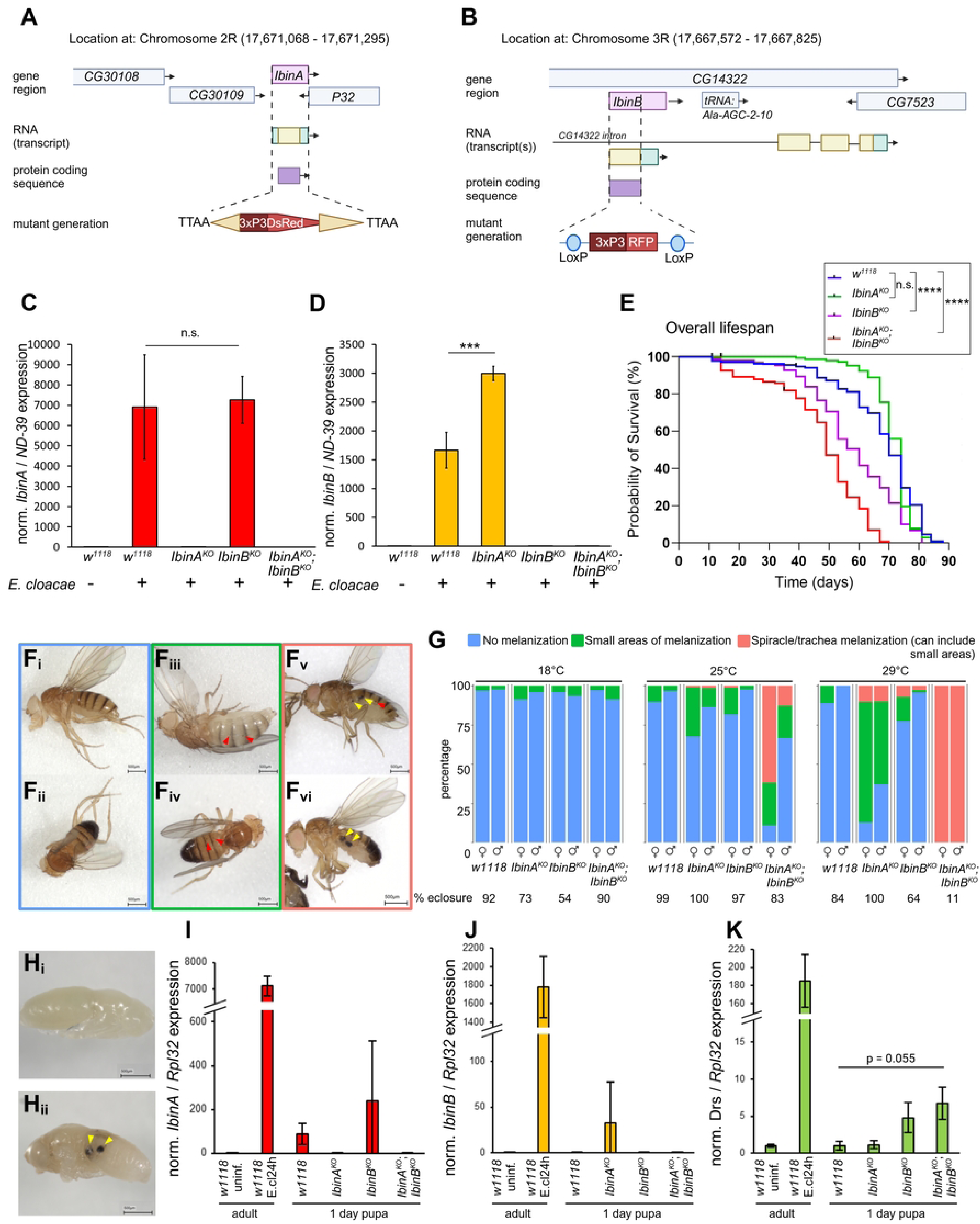
Flies lacking *IbinA*, *IbinB*, or both genes show differences in lifespan and the presence of melanized tissue in adult flies. Schematic representation of the deletion of **A)** the *IbinA* gene (resulting in *IbinA^KO^*), with replacement by the 3xP3DsRed cassette and **B)** the *IbinB* gene (resulting in *IbinB^KO^*), with replacement by the 3xP3RFP cassette. C) *IbinA* expression is not detected in the *IbinA^KO^* and *IbinA^KO^*; *IbinB^KO^* lines when infected with *E. cloacae*, but it is induced normally in *IbinB^KO^* flies. D) *IbinB* expression is not detected in *IbinB^KO^* or *IbinA^KO^*; *IbinB^KO^* flies when infected with *E. cloacae*, but it is expressed more in *IbinA^KO^* flies compared to wildtype. **E)** Overall lifespan of control and mutant flies. n = 150 flies per line. **F)** Representative images of flies with melanized tissue, indicated by arrows, corresponding to categories in panel G: **i)** female without melanized tissue, **ii)** male without melanized tissue, **iii)** female with small areas of melanization, **iv)** male with small areas of melanization, **v)** female with large melanized spots (spiracle/trachea melanization) and **vi)** male with large melanized spots (spiracle/trachea melanization). **G)** Proportions of the melanization phenotype in control and mutant lines in flies reared at 18°C, 25°C and 29°C. **H)** Representative images showing **i)** a healthy control and **ii)** a double mutant (*IbinA^KO^*; *IbinB^KO^*) pupa with melanized tissue. Expression of I) *IbinA*, J) *IbinB* and K) *Drs* in one-day-old pupae of wildtype and mutant flies compared to uninfected and infected (*E. cloacae* 24 hpi) adults. **C-E)** Statistically significant differences are marked with stars: * = p<0.05; ** = p<0.01; *** = p<0.001, **** = p<0.0001.

As melanotic nodules are often formed by (aberrant) hemocyte activation at the larval stage (35), we checked hemocytes of *IbinA^KO^, IbinB^KO^* and the double mutant larvae, as well as control larvae, by classifying hemocytes as baseline immune cells (plasmatocytes, present in a healthy larvae) and immune-inducible, activated hemocytes (mainly lamellocytes, but also other irregularly shaped hemocytes), based on the cell size and shape (36,37) (S5B_i_-B_iii_ Fig). Plasmatocyte and activated hemocyte numbers in both uninfected and parasitoid wasp-infected larvae were somewhat elevated in the *IbinB^KO^* and *IbinA^KO^*; *IbinB^KO^* larvae compared to control and *IbinA^KO^* larvae (S5C_i_-S5C_iv_ Fig). This increase could be due to enhanced release of hemocytes from sessile compartments or from the lymph gland, or increased proliferation in *IbinB^KO^* larvae, all of which are known to occur as a response to parasitoid wasp infection (38). However, as the increase in cell numbers in uninfected larvae is rather small and we do not observe the melanotic nodules at the larval stage, this is unlikely to be the cause of the melanotic masses in the *IbinA^KO^*; *IbinB^KO^* flies.

Many AMPs have been shown to be upregulated during insect metamorphosis (39–42). To see if *Ibin* genes show similar expression, we performed RT-qPCR on approximately one-day-old pupae from wildtype and all three mutant lines. This confirmed that *IbinA* is expressed at a heightened level during the early pupal stage compared to expression in adult flies (Fig 3I), in line with published high throughput expression data (42,43). We were, conversely, not able to show elevated *IbinB* expression at this stage (Fig 3J). Our RT-qPCR results also show increased *Drs* expression in *IbinB^KO^* and *IbinA^KO^;IbinB^KO^* flies (Fig 3K), suggesting possibly dysregulated humoral immunity during pupariation in mutant flies as a contributor to the melanization phenotype that we observe. The full explanation for the requirement of both *IbinA* and *IbinB* in this developmental stage requires further investigation.

### *IbinA* and *IbinB* affect susceptibility to bacterial infections

The upregulation of *IbinA* and *IbinB* transcripts in infected flies (21) (Fig 1) suggests a role for these peptides in the immune response, but they likely do not have a direct bactericidal role (Fig 2). To further investigate their involvement in the immune response, infection experiments were carried out with adult male flies infected with selected bacteria. The wounding injury without any pathogenic bacteria was tolerated normally by *IbinA^KO^*, *IbinB^KO^* and the double mutants (Fig 4A). Infection with *E. cloacae* showed overall very little pathogenicity, although *IbinA^KO^* and *IbinA^KO^*; *IbinB^KO^* flies showed slightly higher rates of dying (Fig 4B). Feeding flies with the pathogen *Serratia marcescens* resulted in clear differences in survival, with *IbinA^KO^* flies surviving best, and all mutant lines showing improved survival compared to the wildtype (Fig 4C).

**Figure 4.**
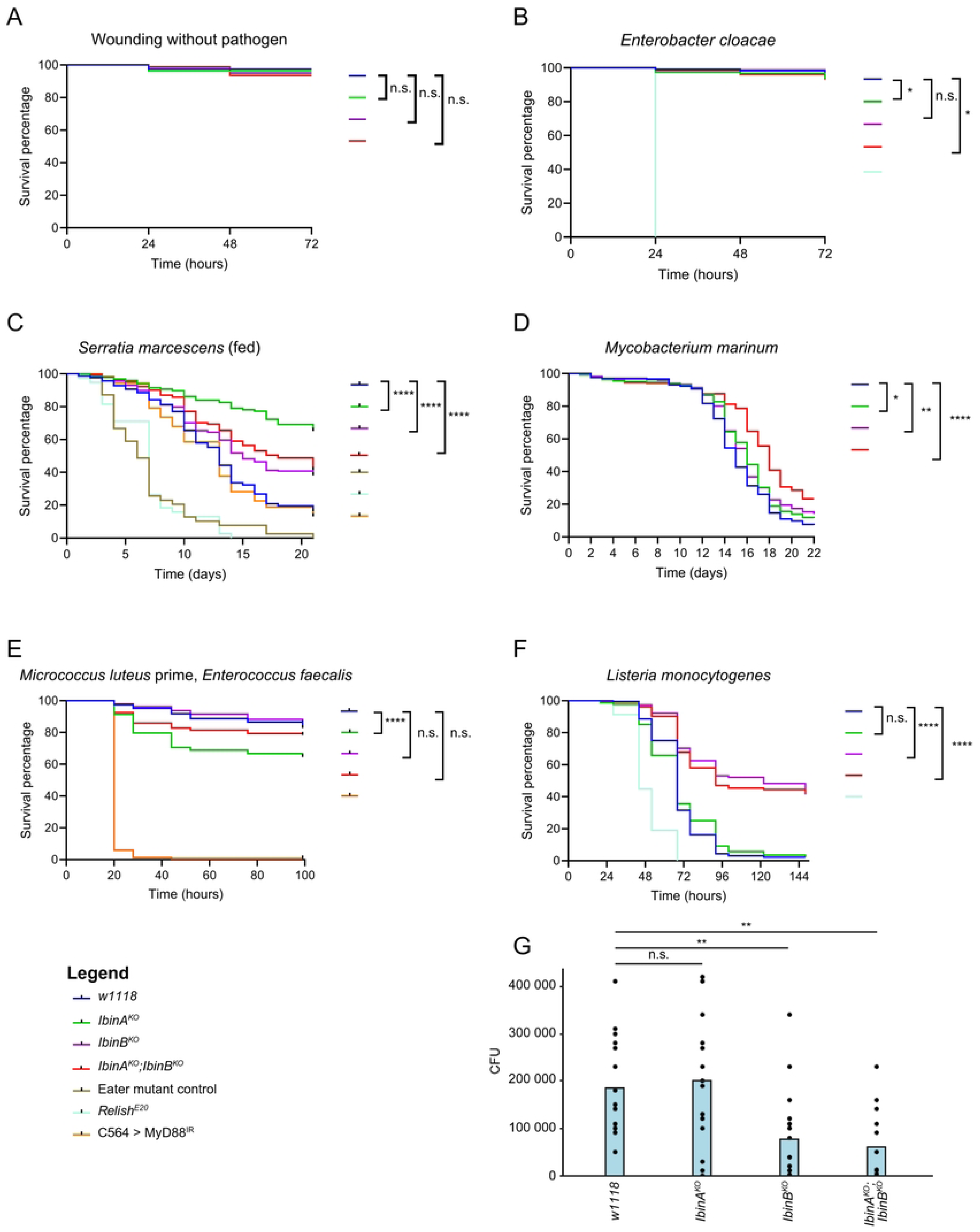
***IbinA* and *IbinB* mutants have distinct roles in immunity**. **A)** *IbinA* and *IbinB* mutants recover normally from wounding without a pathogen. **B, D-F)** Survival of flies after infection by septic injury with selected extracellular and intracellular bacteria. **C)** Survival of mutant and wildtype flies after feeding with *S. marcescens* bacteria. **B-F)** *Relish^E20^* (deficient in Imd pathway response), *eater* mutant (deficient in phagocytosis) and *MyD88* RNAi (deficient in Toll pathway response) flies were used as positive controls. Survival experiments are combined results from three individual experiments. **G)** Bacterial loads in individual flies 36 hours after *L. monocytogenes* septic injury. **A-F)** Results represent three replicates pooled, with 80 individuals per line, per replicate. **A-G)** Statistically significant differences are marked with stars: * = p<0.05; ** = p<0.01; *** = p<0.001, **** = p<0.0001.

Next, we infected flies with the intracellular bacterium *Mycobacterium marinum*, which, similarly to *Mycobacterium tuberculosis* in humans, infects phagocytic immune cells (44). Double mutant (*IbinA^KO^*; *IbinB^KO^*) flies survived the *M. marinum* infection longer than the wild-type controls, and a slight improvement was seen with the single *IbinA* and *IbinB* mutants (Fig 4D). To study the role of the Toll pathway, we first infected flies with *M. luteus* in order to activate the Toll pathway, followed by *E. faecalis* infection 24h later, as in (45). In this experiment there was a significant reduction of survival in *IbinA^KO^* flies compared to wildtype flies and the other mutant lines (Fig 4E). We then tested *Listeria monocytogenes*, known to cause intracellular infection. Infecting flies with *L. monocytogenes* showed a clear difference in survival with flies lacking *IbinB* showing better survival compared to control flies and *IbinA^KO^* flies (Fig 4F). The double mutant flies reproduced the phenotype of *IbinB^KO^* individuals, being more resistant to *L. monocytogenes* infection than *IbinA^KO^* flies and *w^1118^* controls.

To gain a better understanding of the processes behind the difference in the *L. monocytogenes* survival between the *IbinA* and *IbinB* mutants, we investigated whether the difference is due to resistance or tolerance to the bacteria. We infected wildtype and mutant flies with *L. monocytogenes* and homogenized the flies at 36 hours post infection. Plating the homogenates and counting colony forming units shows a significant reduction in the numbers of bacteria in the *IbinB* deficient flies at this timepoint (Fig 4G), suggesting that flies lacking *IbinB* have enhanced resistance, as they can clear bacteria more effectively, or inhibit their growth.

### *IbinA^KO^* and *IbinB^KO^* have distinct effects on gene expression upon *L. monocytogenes* infection

With the aim of gaining insight into the mechanisms behind the different survival phenotypes between *IbinA* and *IbinB* mutants, we infected single and double mutants as well as wild-type controls with *L. monocytogenes* and collected samples at the 36-hour timepoint for RNA sequencing. We also included uninfected samples from all four lines for comparison. Principal component analysis (PCA) of the RNA sequencing results for infected samples shows clustering of *IbinB^KO^* and *IbinA^KO^;IbinB^KO^* samples, separated from *w1118* and *IbinA^KO^* samples by principal component 2 (Fig 5A), consistent with the infection survival results. PC2, however, only accounts for 3% of the total variation between samples. Figure S6A shows the PCA of uninfected samples, with all samples in close proximity along the PC1 axis, which accounts for 99% of variation. This fits our understanding of *IbinA* and *IbinB* genes showing extremely low expression in unchallenged flies, and therefore unlikely to have significant biological effects in this context.

**Figure 5.**
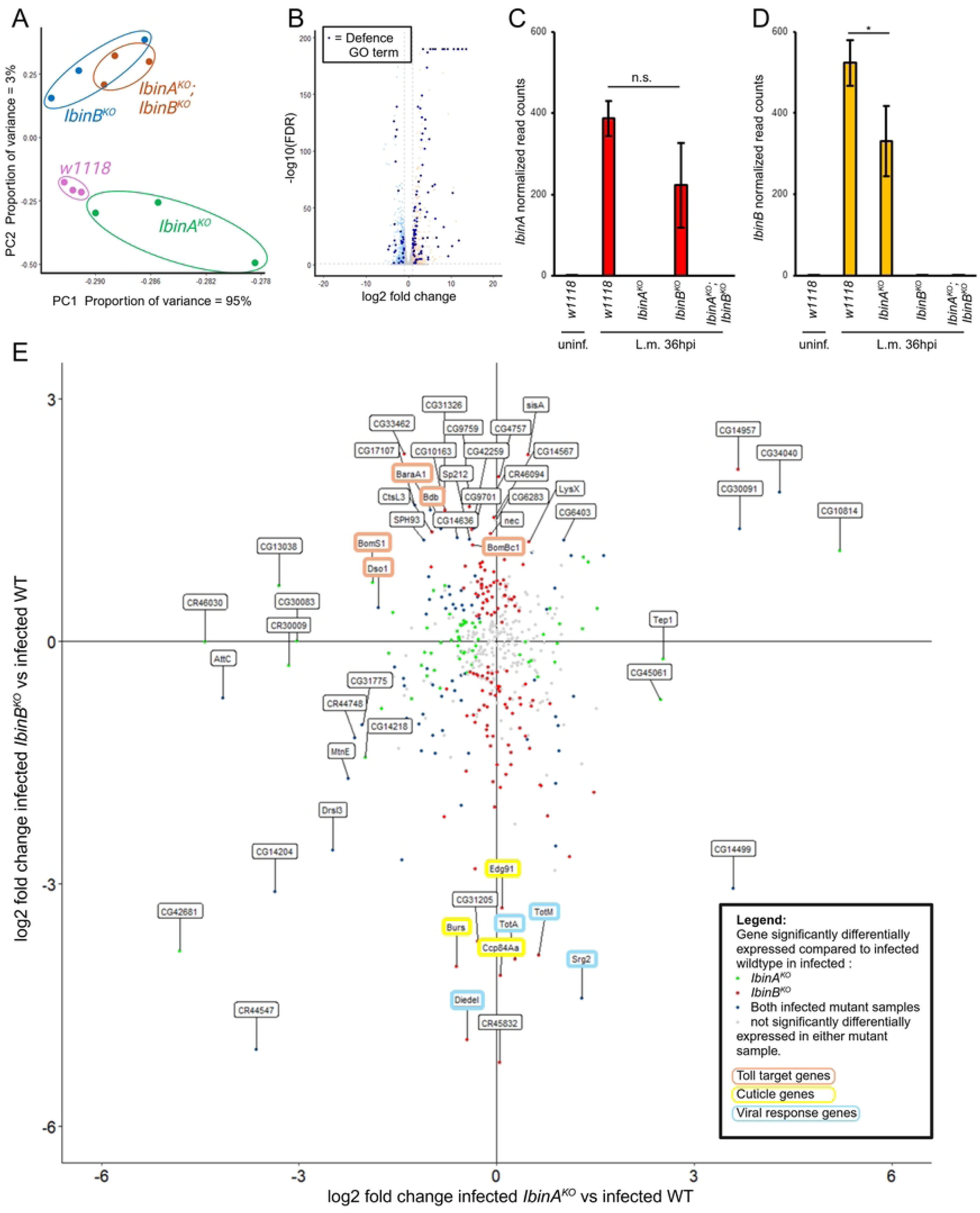
Transcriptional response to *L. monocytogenes* infection in *IbinA* and *IbinB* mutant flies. **A)** PCA plot of infected samples. **B)** Summary of the transcriptional response of wildtype flies to *L. monocytogenes* infection. Genes belonging to the GO term ‘defence response’ are highlighted in blue. RNA sequencing data for **C)** *IbinA* and **D)** *IbinB* expression in *L. monocytogenes* infected flies, normalized to wildtype uninfected flies. Statistically significant difference is marked with a star: * = p<0.05. **E)** Genes with significantly increased expression in wildtype infected flies vs. wildtype uninfected flies plotted by expression pattern in infected *IbinA^KO^* (x axis) or *IbinB^KO^* (y axis) flies compared to infected wild type. Green dots signify significantly differing expression in infected *IbinA^KO^* flies compared to wild type. Red dots signify significantly differing expression in infected *IbinB^KO^* flies compared to wild type. Blue dots signify significant difference in both mutant lines. Highlighted gene labels show increased expression of several Toll pathway targets in *IbinB^KO^* flies, and reduced expression of several viral defense and cuticle proteins in this mutant.

Lists of gene ontology (GO) terms for groups of differentially regulated genes between treatments were produced by filtering differentially regulated genes for up-or downregulation greater than two-fold, and an adjusted P value less than 0.1. Using these criteria, in wildtype flies *L. monocytogenes* infection resulted in the upregulation of 476 genes (S2 Table), and the downregulation of 585 genes (S3 Table). For upregulated genes, top GO terms included ‘response to biotic stimulus’, as well as ‘defense response’ (S4 Table). Top GO terms from the list of downregulated genes were focused on metabolic processes (S5 Table). Highlighting genes in the ‘defense response’ GO term among all differentially regulated genes shows defense response genes both up and downregulated. However, the majority of these genes are upregulated with *L. monocytogenes* infection and make up most of the most highly upregulated genes (Fig 5B).

We began our analysis of gene expression in the mutant lines by examining *IbinA* and *IbinB* levels. Both *IbinA* and *IbinB* are highly upregulated by *Listeria* infection (Fig 5C, 5D), whereas in the mutant lines their expression is absent in both uninfected and infected flies (Fig 5C, 5D). In *Listeria*-infected flies, *IbinA* expression was on average lower in *IbinB^KO^* flies than in wild-type controls, but this difference was not statistically significant (Fig 5C). However, *IbinB* expression is significantly reduced in *IbinA^KO^* samples compared to wildtype flies (Fig 5D).

Following our initial analysis, we looked in detail at the differences in gene expression in the infected fly lines. To examine the differences in response to infection between the mutant lines, we plotted the 476 genes upregulated by *L. monocytogenes* infection in the wildtype flies by expression fold change in single-mutant flies compared to the wildtype (Fig 5E). Most genes clustered around the center of the graph, signifying no difference in expression in mutants compared to the wildtype. Colored points indicate significant difference (compared to the wildtype) in expression of the gene in flies lacking *IbinA* (green dots), *IbinB* (red dots) or in both cases (blue dots). Genes more highly expressed in *IbinB^KO^* included several Toll pathway effectors, namely *BaraA1*, *BomS1*, *Dso1*, *Bdb*, and *BomBc1*, as well as serine proteases such as *Sp212*, *nec*, and *SPH93*. Genes with reduced expression specifically with *IbinB* absent included *Edg91*, *Burs*, and *Ccp84Aa,* all known to have roles in cuticle formation and pigmentation (46–48), and *Diedel*, *Srg2*, and the turandots *TotM* and *TotA*, with roles in viral defense.

In Figure 6, differences in expression of selected immune-relevant genes between the mutant lines and the wildtype are shown. Imd target genes (49) show little significant changes in expression in *IbinA^KO^* flies, but many of these genes show significantly reduced expression in *IbinB^KO^* and double mutant flies (Fig 6A). This includes *edin*, a gene regulating hemocyte numbers during wasp infection (50), and several members of the Attacin, Cecropin, and Diptericin families. In contrast, several Toll pathway-regulated genes (49,51,52) show significant downregulation in *IbinA^KO^* flies, and significant upregulation with *IbinB^KO^* and in double mutant flies (Fig 6B). Genes regulated in this way include *Drs, Bbd*, and *BaraA1*. Several other genes (including many Bomanins) showed the same trends in expression, but lacking statistical significance in either *IbinA^KO^* or *IbinB^KO^* flies. We confirmed this pattern by RT-qPCR of *Drs* expression at 12-and 36-hour timepoints after *L. monocytogenes* infection, showing that *Drs* expression is particularly highly elevated in flies lacking *IbinB* at 36 hours after infection (S7A Fig).

**Figure 6.**
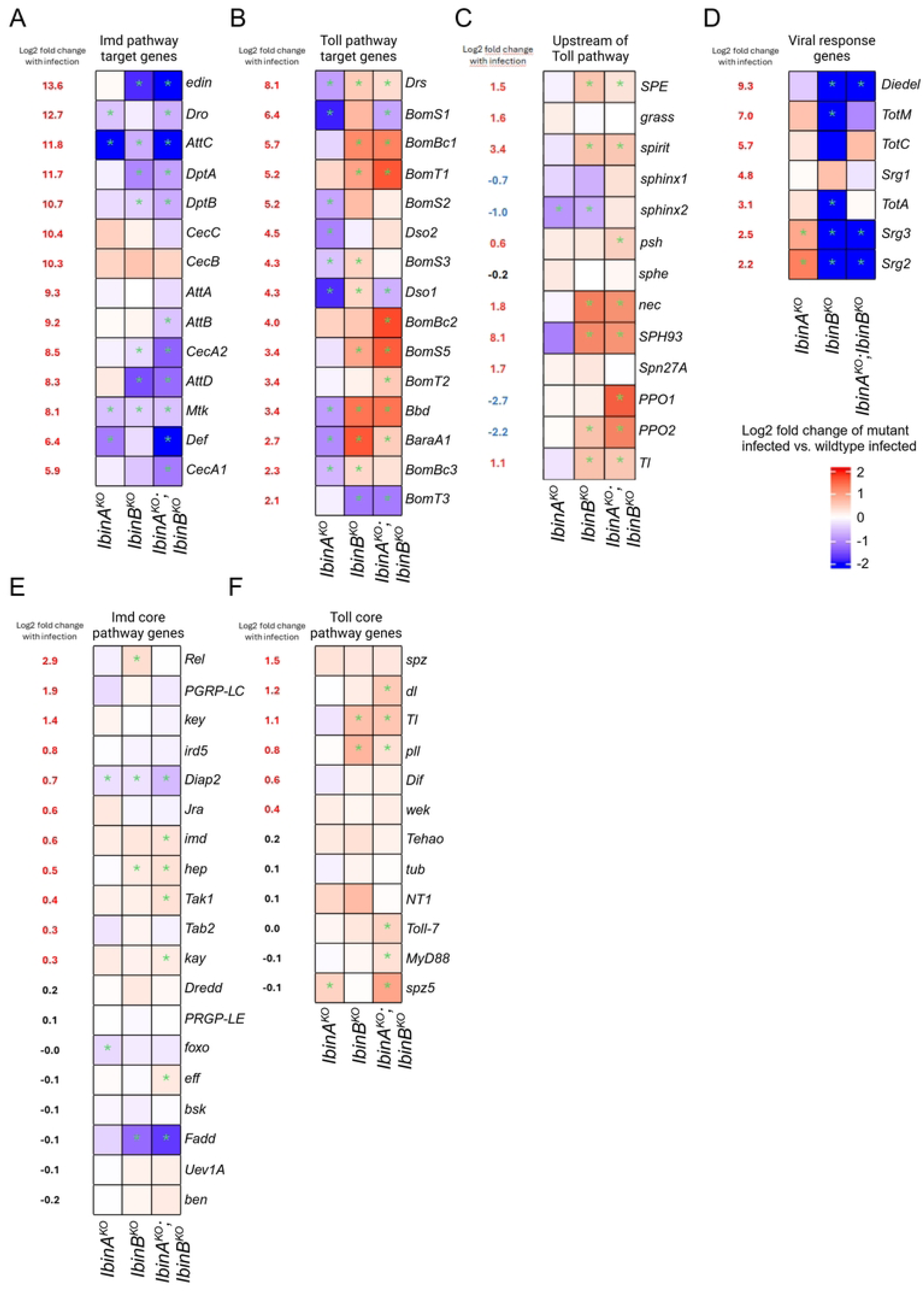
*IbinA^KO^* and *IbinB^KO^* flies have distinct transcriptional responses to *L. monocytogenes* infection. Heatmaps of genes selected for their known roles in the immune response. Leftmost column shows change in expression with infection in wildtype flies, with the following columns comparing infected mutant flies to infected wildtype. **A)** Imd-responsive antimicrobial peptide genes. **B)** Toll pathway effector genes with Log2 fold change > 2 in *L. monocytogenes*-infected wild type compared to uninfected wild type. **C)** Components of the Toll pathway upstream of the Toll receptor. **D)** Genes involved in the response to viral infection with Log2 fold change > 2 in *L. monocytogenes*-infected wild type compared to uninfected wild type. **A-F)** Asterisks signify adjusted p-value < 0.05.

Expression analysis of genes upstream of the Toll receptor in the Toll pathway, and genes in the melanization cascade, shows a similar pattern to that seen in Toll pathway effectors (Fig 6C). This list includes several serine proteases, many of which are also upregulated with infection. The serine proteases *SPE*, *spirit*, *psh*, and *SPH93* all show increased expression in *IbinB^KO^* and double mutant flies. The Toll receptor itself, the negative regulator *nec*, as well as *PPO1* and *PPO2* also show this pattern. In conclusion, these data suggest that IbinA and IbinB have distinct effects on Toll pathway target gene expression upon a challenge with the intracellular pathogen *L. monocytogenes*.

As some genes contributing to viral defense were specifically downregulated in *IbinB^KO^* flies (Fig 5E), and may also affect defense against intracellular bacteria, we took a closer look at their expression (Fig 6D). Sting-regulated genes *Srg2* and *Srg3* show a clear reduction in expression in *IbinB^KO^* and double mutant flies, whereas they have increased expression in *IbinA^KO^* flies. Turandots *TotM*, *TotC*, and *TotA* are all highly upregulated with *L. monocytogenes* infection in our RNA sequencing data (Fig. 6D). *TotM* and *TotA* both show significantly reduced expression in *IbinB^KO^* flies compared to the wildtype. We do not see this pattern in *IbinA^KO^;IbinB^KO^* flies (Fig 6D), making turandots unlikely to be key contributors to the survival changes we see in *Listeria*-infected flies. Of note, *Diedel* shows a very clear reduction in expression in *IbinB^KO^* and double mutant flies. Previous work has identified *Diedel* as a negative regulator of immunity, and specifically of the IMD pathway (53). To this end we investigated a possible role for *Diedel* in septic *L. monocytogenes* infection. However, *Diedel* mutant flies were highly susceptible to *L. monocytogenes* infection (S7C Fig), and thus *Diedel* does not explain the increased resistance of *IbinB* mutants.

In addition, we looked at genes central to the Imd (Fig 6E) and Toll (Fig 6F) signaling pathways, and scavenger receptors (Fig S6D). Imd pathway core components showed no clear pattern, likely reflecting their generally stable expression, including during an immune response. Toll pathway core genes (Fig 6F) showed a pattern similar to the pattern observed in genes in the pathway upstream of the Toll receptor (Fig 6C), and in the Toll pathway targets (Fig 6B), with increased expression of some genes in *IbinB^KO^* and double mutant lines. Our list of scavenger receptors does not show a clearly identifiable pattern across the mutants (S6D Fig). Interestingly, *eater* expression is highly elevated in *IbinB^KO^* and double mutant flies. We verified this pattern of *eater* expression in *Listeria*-infected flies at 12-and 36-hour timepoints (S7B Fig). *Eater* mutant flies show decreased survival time after *Listeria* infection (S7D Fig), and thus this change in *eater* expression may, in part, explain the enhanced resistance of *IbinB^KO^* and double mutant flies.

To investigate the possible involvement of *Ibin* genes in regulating viral infection-responsive genes, we examined the expression of genes responsive to 2’3’-cGAMP injection (signifying STING pathway regulation), using lists of genes from the work of Hédelin and colleagues (54) (S6B, S6C Fig). *L. monocytogenes* infection results in similar regulation of many of these genes, i.e., genes that are upregulated after 2’3’-cGAMP injection are also upregulated in response to *L. monocytogenes* infection (S6B Fig), and *vice versa* for downregulated genes (S6C Fig). Many genes upregulated by 2’3’-cGAMP injection show increased expression in *IbinA^KO^* flies and reduced expression in flies lacking *IbinB* (S6B Fig).

Overall, our RNA sequencing analysis shows *IbinB^KO^* and double mutant flies with increased expression of the Toll-regulated genes including the genes regulating the Toll pathway activity. Increased expression of Toll pathway target genes may explain, in part, the enhanced resistance of *IbinB* mutants against *L. monocytogenes* infection. Based on the transcriptional analysis we conclude that the IbinB peptide functions in downregulating Toll pathway-regulated genes, including those encoding effector molecules. However, the precise point in the Toll pathway where *IbinB* acts remains to be determined.

### IbinB downregulates Toll pathway genes when flies are infected with fungal pathogens

To study further the role of IbinA and IbinB in Toll pathway-mediated immunity, we performed fungal infection experiments, due to the central role of the Toll pathway in resistance to these pathogens (55). We carried out infections with *Beauveria bassiana*, both through the cuticle (‘natural infection’), and as systemic infections by inoculation of the fly thorax using a needle. Infections with *B. bassiana R444* show *IbinB^KO^* flies surviving significantly better than wildtype flies. *IbinA^KO^* flies, in contrast, were more susceptible. This is the case with both natural (Fig 7A) and septic infection experiments (Fig 7B). RT-qPCR confirmed that both *IbinA* (Fig 7C) and *IbinB* (Fig 7D) are expressed when flies are infected with *B. bassiana*, although *IbinB* upregulation showed a high degree of variation between samples, and therefore did not reach statistical significance. We then measured *Drs* expression at 24 (Fig 7E) and 48 (Fig 7F) hours post-infection with *B. bassiana*, confirming that also with this infection, *IbinB* mutants exhibit significantly higher Toll pathway target gene expression than wildtype flies. These results support our finding that *IbinA* and *IbinB* expression affects Toll pathway target gene expression, and that the *Drs* expression level is correlated to survival rate.

**Figure 7.**
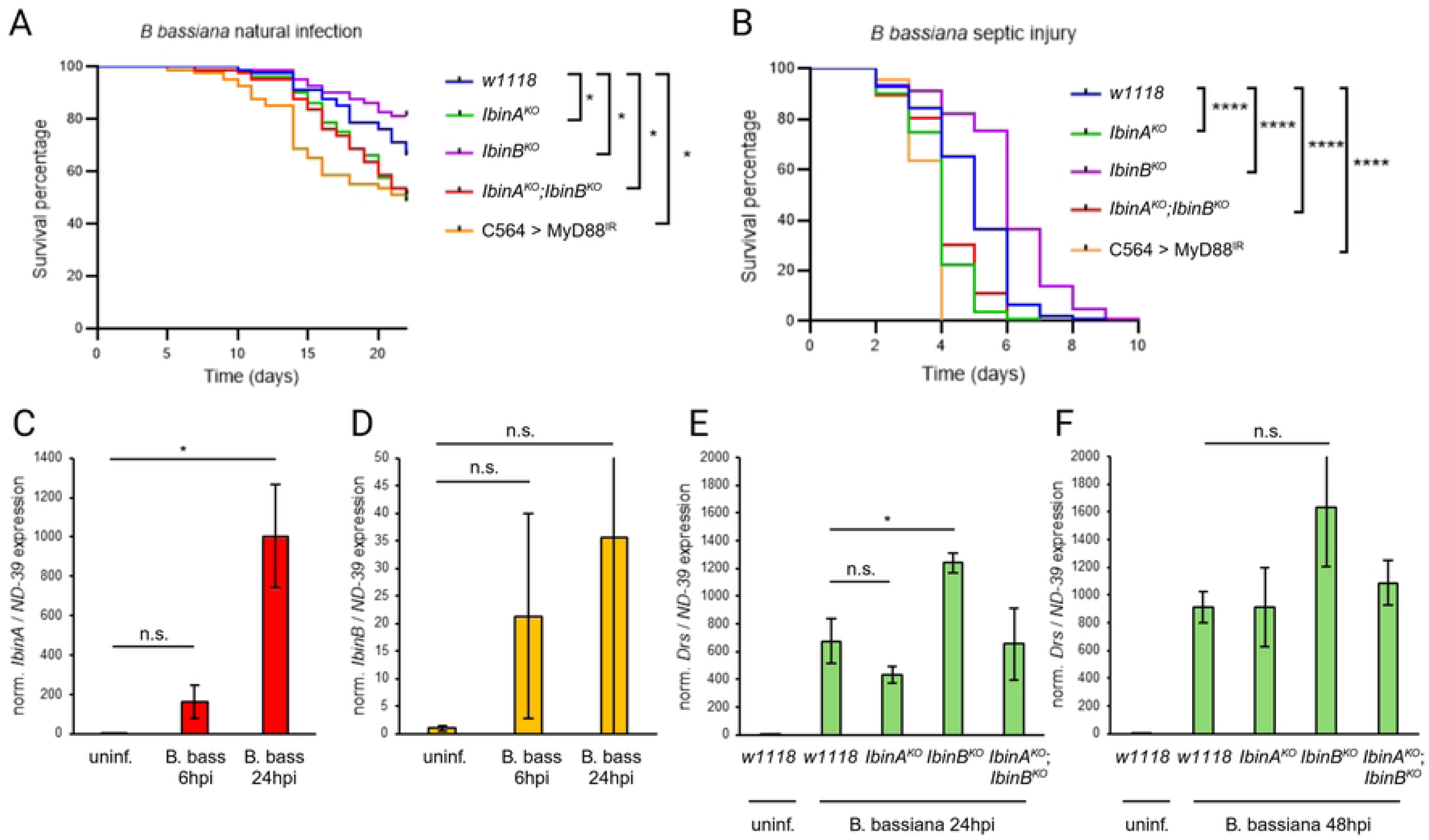
*IbinB* mutants have better survival against fungal infection and higher *Drosomycin* expression. Survival of wildtype and mutant fly lines following infection with *B. bassiana R444* **A)** through the cuticle (so-called natural infection) or **B)** via septic injury. Flies with *MyD88* knockdown in the fat body (*C564*>*MyD88^IR^*) were used as Toll pathway-deficient positive controls. **C)** *IbinA* and ***D)*** *IbinB* expression in wild type flies following septic injury with *B. bassiana*. *Drosomycin* expression was measured **E)** 24 hours and **F)** 48 hours after septic injury with *B. bassiana* from wild type flies and *IbinA^KO^*, *IbinB^KO^*, and double mutant flies. **A, B)** Results represent three replicates pooled, with 80 individuals per line, per replicate. **A-F)** Statistically significant differences are marked with asterisks: * = p<0.05; ** = p<0.01; *** = p<0.001, **** = p<0.0001.

## Discussion

Our previous study identified *IbinA* and *IbinB* as among the most highly upregulated immune responsive genes in *Drosophila* in a wide range of infection contexts (bacterial infections as well as wasp parasitization) (21). In this work we establish these two genes as being evolutionarily conserved among related fly species, suggesting an important and established role in the host immune response in these insects. Despite their sequence similarity, our infection results and RNA sequencing data highlight differing roles for these peptides. We add to the range of bacterial pathogens studied and show that fungal infection also triggers expression of both *IbinA* and *IbinB*.

The Ibin peptides appear to be unique for the subgenus *Sophophora*. Such evolutionary novelties are not rare among genes involved in immunity, and their origin is an important question. The replacement of the more ancestral Mibin with Ibin suggests a sudden shift in function of the gene in the ancestor of all sophophorans, perhaps because of a corresponding change in the target of these peptides. This brings us to the possible origin of Mibins in the first place. We consider it noteworthy that the Ibin genes are generally located near the *Diptericin*/*Attacin* loci in the genome of *D. melanogaster*. Many attacins and diptericins have a proline-rich N-terminal domain, and it has been suggested that this domain may have a common origin with proline-rich antimicrobial peptides such as Drosocin, and perhaps also Metchnikowin (29). This idea has been further extended to include the Ibins as well as other proline-rich peptides (56). In calyptrate flies, such as *Lucilia*, *Cochliomyia* and *Sarcophaga*, the *Mibin* and *Attacin A*-like genes are even situated just next to each other (S2 Fig). *IbinA* and *IbinB* share a pattern of transcription common among immune regulated genes, with extremely low baseline levels in uninfected flies, apart from the higher expression observed during pupariation. These genes show upregulation of several hundred-or thousand-fold upon infection (with *IbinA* and *IbinB* among the most highly upregulated genes during infection). Most genes with such transcription patterns are examples of the classic AMPs, such as Cecropins, Attacins, and Diptericins, normally described as Imd-regulated, and the generally Toll pathway-regulated Drosomycin and Bomanins. As effector molecules, large quantities of these peptides are produced to combat infection. Similarly regulated genes also include cytokines such as the Unpaired (Upd) genes, and several serine proteases and serine protease homologues, many of which have known roles in the Toll and melanization pathways, with others having unknown or poorly defined roles in the immune response (57). Stress-related genes, including the Turandots, are also frequently among the most highly upregulated genes with infection. Rommelaere and coworkers (58) recently presented evidence that Turandots play a tissue protective role in the context of the immune response.

Our results provide insight into where *IbinA* and *IbinB* fit within this transcriptional context. We did not find any direct bactericidal or bacteriostatic effect of Ibin peptides, pointing away from roles as AMPs. We also did not find any increased expression with heat shock. We replicated previous results showing upregulation of *IbinA* in certain stress conditions (visual exposure to parasitoid wasps, and social isolation of male flies) (23,24). This upregulation is variable between individuals, and at a much lower level than in the case of infection. In summary, there is some evidence of a role for *IbinA* (as with other immune induced genes) in the transcriptional response to some stress conditions. However, the physiological consequences of this response require further investigation.

While *Ibin* genes do not have significant roles as AMPs or in stress response, our results do suggest an immune regulatory role for *IbinA* and *IbinB.* The RNA sequencing results show that during *L. monocytogenes* infection, flies lacking *IbinB* had increased expression of Toll pathway-mediated effectors, including most Bomanins and *Drs*. In contrast, *IbinA^KO^* flies had reduced expression of many of these genes. This fits the pattern we observed across infection experiments, with *IbinB^KO^* flies surviving longer in infections where the Toll pathway plays a significant role in the immune response, such as *E. faecalis* infection, as well as infection with fungal pathogens. Both Toll and Imd pathways have been shown to be important in survival after *L. monocytogenes* infection (59). These infection and transcriptome results lead us to suggest that IbinB acts as a negative regulator of Toll pathway target gene expression, with IbinA having the opposite role. The overall lifespan of the mutant lines also supports this conclusion; flies lacking *IbinB* have shorter overall lifespan, which could be due to overactivation of the Toll pathway causing tissue damage. While a more robust Toll pathway response increases the ability to clear many pathogens, immune dysregulation has detrimental effects throughout the animal’s lifespan. We also show a dependence of *IbinB* expression on the Imd pathway, including when flies are infected with bacteria expected to trigger a Toll pathway-mediated response. This finding remains to be explained, perhaps providing evidence for crosstalk and mutual regulation between Toll and Imd pathways.

The STING pathway has been shown to be involved in the response to *L. monocytogenes* infection in *D. melanogaster* (60). In our results, many STING/cGAS regulated genes are differentially regulated in flies lacking *IbinB*, in most cases downregulated. Overall, these RNA sequencing results suggest a role for *IbinB* in modulating different aspects of the immune response, increasing the antiviral transcriptional response and reducing the expression of the Toll-responsive effector genes.

The infection survival results and transcriptome data do include findings that cannot be clearly explained by regulation of the Toll pathway. During *S. marcescens* infection *IbinA^KO^* flies exhibit the best survival of all lines, followed by *IbinA^KO^;IbinB^KO^* and *IbinB^KO^*. The Toll pathway does not play a significant role in resistance to this bacterial species, suggesting that a different explanation is needed for these results. Scavenger receptors were differentially regulated in *Ibin* mutants, in particular *eater*. This perhaps suggests changes in the numbers or activation state of hemocytes in flies lacking *Ibin* genes. This is consistent with our findings of changes in hemocyte numbers and activation state in larvae in these mutant lines, although how the larval hemocyte state translates to immunity in adults is not clear. The cellular immune response has previously been shown to be significant in resistance to *S. marcescens* in *Drosophila* (61).

The blackened tissue we observed in adult mutant flies, likely around parts of the tracheal system of the flies, indicates that in the special pupal immune environment, Ibin peptides play a protective role, either by directly protecting tissue from attack by AMPs (as reported for Turandots (58)), or by regulating the melanization response. Unlike Turandots, however, we were not able to show upregulation of *Ibin* genes in response to heat or osmotic stress.

Tang and coworkers (62) observed melanization of the trachea of flies (already in larval stages) lacking the serpin Spn77Ba. This melanization led to local and systemic expression of *Drs* via the Toll pathway. The melanization we observed appears during pupariation and early adulthood, and it generally does not cover the whole trachea. However, the melanization phenotype, combined with the increased expression of Toll pathway targets, including *Drs*, points towards a role for IbinA and IbinB in regulation of serine protease activity, and thus both melanization and Toll pathway activity. The process that occurs at pupariation that leads to aberrant melanization in the mutant flies remains to be explored, as does the reason for the increased penetrance of the melanization phenotype as rearing temperature increases, and the larger melanotic spots observed at the trachea, predominantly of female flies. Flies express many AMPs at pupariation, suggesting that negative regulation of the immune system might be particularly important at this stage to prevent either aberrant melanization or tissue damage caused by humoral immune effectors. Our data suggest that *Ibin* genes are mainly expressed during pupariation and when flies are infected, providing further evidence that they are required to modulate the humoral immune response and/or melanization under specific circumstances. Scherfer et al. (34) also showed some melanization of the trachea in flies lacking proper negative regulation of melanization, suggesting that the trachea is particularly susceptible to aberrant melanization. They speculate that this is due to oxygen being a substrate for phenoloxidase (34).

Overall, our results uncover roles for IbinA and IbinB in regulating the immune response. Specifically, IbinB has a negative regulatory effect on the expression of Toll pathway effector genes while IbinA in the context of infection appears to play the opposite role. In the case of *L. monocytogenes* infection, as well as resistance to *E*.

*faecalis* and fungal pathogens, *IbinB* mutants are able to mount a more effective response, likely due to their higher expression of Toll effectors, including *Drs*. While *IbinB^KO^* flies consistently survive better than wild type individuals in the context of bacterial infection, these flies have overall shorter lifespans, which is further evidence for the role of *IbinB* in regulating the immune system, thus preventing excessive tissue damage from aberrant immune activation. Observations of melanization during pupariation further suggest an immune regulatory and tissue protective role for Ibin genes. Our findings on the roles of *Ibin* genes in immune regulation highlight the importance of appropriate levels of immune activation, both during immune challenge, and in unchallenged conditions throughout development.

## Materials and Methods

### Bioinformatic analysis of *IbinA*, *IbinB* and *Mibin* genes

*Ibin* homologs were initially identified by simple blastp and tblastn searches of the *refseq_protein* and *refseq_genomes* databases, followed up by repeated searches with some of the initially retrieved hits, and similar searches of the *wgs* database for selected taxa. To limit the scope of the study, we have refrained from expanding the set as new species have been added to the databases. A consensus phylogenetic tree, showing the occurrence and chromosomal locations of *IbinA*, *IbinB* and *Mibin* orthologs, was built from (26,63,64), with time calibration from (26). The first *Mibin* homologs were identified by inspecting all open reading frames in the 1 kb intergenic region between *Drosophila busckii* homologs of *D. melanogaster* genes *CG30109* and *P32*, where *IbinA* is located. Further *Mibin* homologs were then identified by tblastn searches of the *refseq_protein*, *refseq_genomes* and *TSA* databases. Multiple sequence alignments were done by Clustal Omega at the The EMBL-EBI Job Dispatcher (https://www.ebi.ac.uk/jdispatcher/msa/clustalo), and further manually curated. Signal peptide sequence prediction was done using SignalP-6.0 at the DTU Health Tech server (https://services.healthtech.dtu.dk/services/SignalP-6.0).

### *In vitro* bacterial killing assay

The *in vitro* bacterial killing assay was modified from (65). Briefly, the selected bacteria were grown overnight either in Luria-Bertani (LB) or brain heart infusion (BHI) medium as detailed above. The next morning, bacteria were subcultured in fresh medium for 1-2 hours to reach OD600 of 0.33 (exponential growth phase). 1:10 dilution of bacteria was added in a fresh solution of LB/BHI including the peptide tested or antimicrobial agent control, at desired final concentrations.

Mature IbinA and IbinB peptides were obtained from GenScript Biotech (Netherlands, B.V.) as lyophilized powder. Peptide sequences are as follows: IbinA: KNHEDWGGYRPSDYDPRPYFRQF and IbinB: KNHEEWKGQRPWDYDRRPPSN PYA. Peptides were dissolved in ultrapure H_2_O and kept at-80°C until use. The following antimicrobial agents were used as positive controls: Cecropin A (Sigma Aldrich, #C6830) for *Enterobacter cloacae* and *Escherichia coli*, Melittin from honeybee venom (Sigma Aldrich #M2272) for *Staphylococcus aureus* and Lysozyme (EMD Millipore Corp., USA, #4403) for *Enterococcus faecalis*. Sterile ultrapure H_2_O was used as the negative control. Samples were incubated for 2h at 25°C, after which they were placed on ice. Serial dilutions of bacteria were carried out in ice-cold 1 x PBS and 3 µl (four replicates per treatment) were plated on LB/BHI agar. Plates were grown overnight at 29°C. Colony forming units (CFU/ml) were counted from the appropriate dilution on the agar plate and calculated taking all dilution steps into account.

### *Drosophila* lines

The *D. melanogaster w^1118^* and *Canton S* lines were used as control lines. The *w^1118^* line has been kept in the laboratories of the authors for over 30 years, and our GAL4-driver lines are backcrossed into this genetic background. The UAS/GAL4 system (66) was used for silencing selected genes in the selected tissue. To study the effect of the Toll and Imd pathways, *UAS-MyD88^IR^* (GD #25399), *UAS-Rel^IR^* (GD #49414), *UAS-PGRP-LC^IR^* (GD #51968) and *UAS-Imd^IR^* (GD #9253) RNAi lines from the Vienna Drosophila Resource Center (VDRC) and *UAS-FADD^IR^* (12297R-1, III) from the NIG-Fly center in Kyoto were used. *C564-GAL4* line, driving strong expression of the UAS-construct in the fat body (67) and also some other tissues, was a gift from Prof. B. Lemaitre (Global Health Institute, Swiss Federal Institute of Technology Lausanne, Switzerland). *Relish^E20^* flies contain a null allele of the *Relish* gene (68). The transheterozygous *eater* mutant flies were obtained by crossing together the Bloomington Stock Center deficiency lines *Df(3R)D605* (stock #823) and *Df(3R)Tl-I* (stock #1911), as in Kocks *et al*. (69). *Diedel* knockout line (generated by imprecise p-element excision described in (70)) and its background control line were a kind gift from Professor Jean-Luc Imler (CNRS, Strasbourg, France).

*IbinA* and *IbinB* knock-out (KO) mutants were generated at Wellgenetics Inc. (Taiwan, www.wellgenetics.com) with CRISPR/Cas9 mediated genome editing by homology dependent repair (HDR). For *IbinA^KO^*, one guide RNA and a dsDNA plasmid donor was used, whereas for *IbinB^KO^*, two guide RNAs and a dsDNA plasmid donor was used. The mutants were backcrossed to the control line used in the experiments (*w^1118^*). The *IbinA^KO^*; *IbinB^KO^* double mutant was generated by crossing the backcrossed mutant lines together, to obtain the genotype: *w^1118^*; *IbinA^KO^*; *IbinB^KO^*.

### Imaging pupal & adult melanization, scoring phenotypes

To score melanization phenotypes, ten male and ten female flies were allowed to lay eggs in a vial at 25°C for 24 hours, before being discarded. Vials were then kept at the experimental temperature. When eclosion began, each individual fly was visually inspected using a dissecting microscope and scored into one of three categories: no melanization visible; small areas of melanization; or spiracle/trachea melanization (flies were included in this category if they featured both spiracle/trachea melanization, as well as smaller areas visible through the dorsal cuticle). This was performed for five days from the beginning of eclosion, after which pupal cases were counted to determine eclosion percentage. Representative images of phenotypes were taken from flies raised at 29°C using a Nikon SMZ475T microscope.

### Larval hemocyte analysis

*Ibin* mutant and the control flies were allowed to mate and lay eggs for 24 hours at 25°C. The eggs were then transferred to 29°C for further development. To analyze the hemocytes after wasp-infection, 2^nd^ instar larvae were exposed to 10 female *Leptopilina boulardi* strain G486 wasps for two hours at RT (22°C), after which the wasps were removed, and the larvae placed back at their rearing temperature until they reached mid-to late 3^rd^ instar larval stage. Then the larvae were gently washed by brushing in water to remove food remnants and dissected in 20 µl of 1% bovine serum albumin (BSA) in 1 x PBS. The sample containing hemocytes from the hemolymph was pipetted into a microtube with 80 µl of 1% BSA in PBS. 30 µl of the sample was run with a BD Accuri C6 flow cytometer (BD Biosciences). In the case of wasp-infected larvae, the presence of wasp larva was visually confirmed prior to sample collection. Each genotype was analyzed in triplicate (3 x 10 larvae), with equal numbers of males and females. Hemocytes were first separated from the debris released when dissecting the larvae based on their location in a forward scatter area (FSC-A) – side scatter area (SSA-A) plot (71) (S4A Fig). Separation between inactivated, “basal” plasmatocytes and activated hemocytes was made based on the protocol described (36,37). Shortly, plasmatocytes are small and round in their inactivated state and can be gated using the forward scatter area vs. forward scatter height plot, where round cells form a population detected at an 45° angle.

Lamellocytes are irregularly shaped, which places them outside of the 45° angle in the forward scatter area *vs.* forward scatter height plot.

### Culturing and processing microbes for infections

*Micrococcus luteus* and *Enterobacter cloacae* were cultured on LB agar plates under antibiotic selection, and *Enterococcus faecalis* was cultured in BHI medium as described previously (21). Bacteria were collected from the plate or pelleted and mixed with glycerol to create a thick paste. *Listeria monocytogenes* was grown overnight in 5ml BHI medium, shaking at 37°C, after which the culture was diluted 1:25 in fresh medium and grown to an OD_600_ of 0.75. 1ml of bacterial suspension was centrifuged at 500 x g for 3 minutes. The supernatant was removed, and the pellet is resuspended in 100ul of 50% glycerol. *Mycobacterium marinum* ATCC 927 was grown at 28°C in 7H9 medium supplemented with ADC enrichment and Tween 80 to achieve OD_600_ value of 0.75. 1ml of bacterial culture was centrifuged for 3 minutes at 10,000 rcf and the supernatant removed. Pellet was resuspended in 100ul of 50% glycerol. *Serratia marcescens* Db11 was grown at 37°C in LB broth to OD_600_ of 1.0. *Beauveria bassiana* strain R444 (BB-PROTEC) commercial spores were produced and provided by Andermatt Biocontrol as fungal spores mixed with talk powder.

### Fly infections and survival monitoring

Fly infections were carried out using multiple infection routes and methods. To cause septic injury, flies were pricked in the thorax area with a sharpened tungsten wire dipped in the thick bacterial paste. Infections with *Serratia marcescens* by feeding were carried out essentially as in (61). Briefly, flies were placed in vials with cotton wool plugs soaked with 5 ml either in a 50 mM sucrose solution or the same sucrose solution inoculated 1:100 dilution of *S. marcescens* bacteria. For causing a natural fungal infection, 30 mg of commercial spores were added to an empty fly vial. Flies were added to the vial and tapped to the bottom of the vial for 30 seconds to cover flies in fungal spores, as described previously (45,51). For systemic fungal infections, 30mg of commercial fungal spores were suspended in 100 ul of 50% glycerol. Flies were pricked in the thorax area with a sharpened tungsten wire dipped in the fungal paste. Depending on the rate of dying to the infection, the survival of flies was recorded either several times a day, daily or every other day. For long survival times, the food was changed twice a week until the end of the experiment.

### Bacterial load

Colony forming unit (CFU) per fly, was measured from flies 36 hours post *Listeria monocytogenes* infection as modified from (37). Eight flies per strain were infected. To kill the cutaneous bacteria, each fly was dipped in 70% ethanol for 30 seconds, after which the fly was placed in 100 μl PBS and homogenized with a plastic pestle. A 10-fold dilution series was prepared in 96-well plates, and 3 μl of each dilution was plated on an LB-agar plate, incubated overnight at 37°C. Colonies were counted under light microscope.

### RNA extraction and RT-qPCR

RNAs were extracted and RT-qPCR (reverse transcriptase quantitative polymerase chain reaction) carried out as described previously (72). Three replicates of five male flies per genotype were collected and snap-frozen on dry ice. TRI reagent (MRC; Thermo Fisher Scientific) was added to the frozen flies, and the flies were homogenized using a micropestle (Thermo Fisher Scientific). Thereafter total RNAs were extracted according to manufacturer’s instructions. RNAs were dissolved in nuclease-free water and the concentrations and purity were measured using the Nano-Drop 2000 equipment (Thermo Scientific). RT-qPCR was carried out from the total RNA samples (∼40 ng/sample) with the iTaq Universal SYBR Green Onestep kit (Bio-Rad, Hercules, CA). A gene encoding a subunit of mitochondrial respiratory complex I, NADH dehydrogenase (ubiquinone) 39 kDa subunit (ND-39), was used as a steadily expressing control gene to normalize differences in RNA amounts between samples. The primers used are listed in S6 Table.

### Stress experiments

Heat shock of flies was performed on three-to-five-day old male flies. Flies were maintained at 25°C until the start of the experiment. Fly vials were transferred to a pre-heated water bath at 36°C for one hour or four hours. Immediately after exposure, flies were anesthetized on a CO_2_ pad, decapitated using a scalpel blade, with heads and bodies of flies separately transferred to-80°C storage in preparation for RNA extraction. Osmotic stress experiment was performed by feeding flies normal fly food with 4% NaCl added. Flies were maintained on food for 8 or 24 hours before flash freezing and RNA extraction. Separately, flies were maintained on 4% NaCl food with blue dye (Brilliant blue, Carbosynth) to confirm that 100% of flies maintained on 4% NaCl food consumed the food. Food consumption was confirmed by checking for the presence of blue dye in the fly guts under a light microscope.

For measuring courtship delay time male and female flies were collected into separate vials within six hours after eclosure and maintained for two days at 25°C. Individual male and female flies were transferred using suction (without anaesthetization or tapping) into separated sides of a mating chamber at 8 am. Mating chambers were kept for one hour at 25°C. The divider was then removed, allowing interaction of the male and female flies. Courtship and mating behavior was filmed using a cell phone (iPhone 5S, Apple, United States) placed in the incubator above the mating chambers. Flies were allowed to interact for one hour before the experiment was ended, with time to copulation for individual pairs noted from the video recording.

Social isolation was performed on two-day-old male adults. Flies were transferred by suction into individual vials. Vials were kept in a 25°C incubator for 3 days, separated by dividers to prevent visual contact between individual flies. Individuals were then anesthetized and decapitated in preparation for RNA extraction. Gene expression levels were measured from heads and bodies separately.

### Overall lifespan

Three-to-five-day old adult males were used for lifespan experiments. Flies were maintained in vials of 25 individuals at 25°C. Flies were transferred to new food vials twice per week, with new deaths recorded on days when the flies were transferred. The experiment continued until all flies were dead.

### Sample preparation for RNA sequencing

Flies were infected with *Listeria monocytogenes* using the same protocol as used for infection survival experiments. Live infected flies and uninfected controls with similar incubation times were collected to-80°C at the 36-hour post-infection timepoint. One biological replicate consisted of five individual flies, with three biological replicates per treatment. RNA was extracted using the TRI reagent protocol described above.

### RNA sequencing

Quality control of isolated RNAs, preparation of the RNA library (including quality control), the whole transcriptome sequencing and data analysis were carried out by Novogene Co. Ltd, UK (www.novogene.com). In short, messenger RNA was purified from the total RNA using poly-T oligo-attached magnetic beads. RNA was fragmented, after which the first strand cDNA synthesis was carried out using random hexamer primers, followed by the second strand cDNA synthesis. Following library construction consisted of end repair, A-tailing, adapter ligation, size selection, amplification and purification. The quality-controlled library preparations were quantified, pooled and sequenced using an Illumina platform. Paired end 150 bp technology was used to sequence the samples. Sequence data analysis first included a quality control, where raw reads were processed by removing reads containing adapter, reads containing poly-N and low-quality reads from raw data. Reads were mapped to the *D. melanogaster* reference genome (Genome ID: ensembl_drosophila_melanogaster_ bdgp6_gca_000001215_4). Gene expression levels for each gene were calculated as FPKM (Fragments Per Kilobase Million).

## Statistical analysis

Statistical analyses of RT-qPCR gene expression results were carried out using a two-tailed t-test for two samples assuming equal variances using R version 4.3.3 (73). Statistical analyses of fly survival from infection and lifespan experiments were carried out using the log-rank (Mantel-Cox) test with Prism version 10.3.1 (GraphPad Software LLC). The data on hemocyte counts were analyzed with R version 4.3.3 (73), using a negative binomial generalized linear model (MASS package; (74)). The least square means (estimated marginal means) were analyzed for multiple comparisons (75), and the Tukey method was used for adjusting the p-value. Heatmaps were generated using the ComplexHeatmaps R package version 2.18.0 (76). Differential gene expression was calculated by Novogene using the R package DESeq2 (77), with the false discovery rate calculated using the Benjamini-Hochberg method.

## Data availability

The transcriptome analysis dataset is available via the Gene Expression Omnibus database, using the GEO accession number GSE293484.

## Acknowledgements

The authors would like to thank Päivi Lillsunde (Tampere University) for technical assistance and Mark Hanson (University of Exeter) for advice regarding fungal infections. This work was funded by Sigrid Jusélius Foundation (to M.R. and T.S.S.), the Tampere tuberculosis foundation (to M.K.M., M.R., L.V. and S.V.), the Finnish Cultural Foundation (to M.K.M.), the Swedish Research Council and the Nils Erik Holmsten Foundation (to D.H.) and the Research Council of Finland projects 3121353367 and 3121322732 (to T.S.S.). The Tampere *Drosophila* Facility, providing resources for fly work, is partially funded by Biocenter Finland.

## Author contributions

Conceptualization, M.K.M., S.V., T.S.S and M.R.; Methodology, M.K.M., S.V., L.V., T.T. and T.S.S.; Investigation, M.K.M., S.V., L.V., P.V., E.H., T.T., A.M., T.S.S. and D.H.; Writing – Original Draft, M.K.M., S.V., L.V., T.S.S. and D.H.; Funding Acquisition, M.K.M., S.V., T.S.S., D.H. and M.R.; Resources, S.V., T.S.S., D.H. and M.R.; Supervision, S.V., T.S.S. and M.R., Project administration, T.S.S., D.H. and M.R.

## Supporting information captions

**S1 Figure. Ibin open reading frames.**

**S2 Figure. Chromosomal location of *Ibin* and *Mibin* genes in different species. S3 Figure. Mibin open reading frames.**

**S4 Figure. *IbinB* expression is dependent on functional Imd pathway. *IbinA***

**shows varied expression upon exposure to various stressors.**

**S5 Figure. Example images of *Ibin* mutant phenotypes. Flow cytometric analysis of larval hemocytes in *Ibin* mutants and controls.**

**S6 Figure. Additional heatmaps of immune-relevant genes at 36hpi with *L. monocytogenes* infection in *Ibin* mutants and controls.**

**S7 Figure. qPCR verification of *eater* and *Drs* expression at 36hpi with *L. monocytogenes* infection. Survival of *eater* and *Diedel* mutant flies from *L. monocytogenes* infection.**

**S1 Table. RNA sequencing fold change comparisons.**

**S2 Table. Genes upregulated with L.m. infection in wildtype (w1118).**

**S3 Table. Genes downregulated with L.m. infection in wildtype (w1118).**

**S4 Table. Gene ontology hits from Sup Table 2 list.**

**S5 Table. Gene ontology hits from Sup Table 3 list.**

**S6 Table. RT-qPCR primers used in this study.**

